# Phytochemical Constituency Profiling and Antimicrobial Activity Screening of Seaweeds Extracts Collected from the Bay of Bengal Sea Coasts

**DOI:** 10.1101/680348

**Authors:** Mahmudul Hasan, Md. Abdus Shukur Imran, Farhana Rumzum Bhuiyan, Sheikh Rashel Ahmed, Parsha Shanzana, Mahmuda Akter Moli, Shakhawat Hossain Foysal, Suma Bala Dabi

**Affiliations:** Department of Pharmaceuticals and Industrial Biotechnology, Sylhet Agricultural University, Sylhet-3100; Department of Botany, University of Chittagong, Chittagong, Chittagong-4331; Department of Plant and Environmental Biotechnology, Sylhet Agricultural University, Sylhet-3100; Department of Molecular Biology and Genetic Engineering, Sylhet Agricultural University, Sylhet-3100; Department of Genetic Engineering and Biotechnology, Shahjalal University of Science and Technology, Sylhet-3114

**Author notes:** **Corresponding Author:** Mahmudul Hasan, *Assistant Professor, Department of Pharmaceuticals and Industrial Biotechnology, Faculty of Biotechnology and Genetic Engineering, Sylhet Agricultural University, Sylhet-3100*.

**Keywords:** Antimicrobial activity, Phytochemicals, Seaweeds, *Hypnea musciformis*, *Enteromorpha intestinalis*

## Abstract

Seaweeds are able to produce a great variety of secondary metabolites that are characterized by a broad spectrum of biological activities. Two seaweeds species, namely *Hypnea musciformis* and *Enteromorpha intestinalis were* studied to evaluate the phytochemical constituency and antimicrobial activities. First of all, crude extracts of both sea weeds were prepared by two different extraction methods (soaking and water bath) using different solvents. Phytochemicals profiling results revealed the presence of bioactive compounds (flavonoids, alkaloids, tannin, saponin and phenols) in both seaweed extracts. Quantification results for ethanolic extracts of *H. musciformis and E. intestinalis* estimated 51 mg and 43 mg tannins in per gram of dried samples and flavonoids contents were found 67 mg and 57 mg/g mg QE/g respectively. Total phenolic contents were determined in terms of gallic acid equivalent (GAE). *H. musciformis* exhibited higher amount of phenolics (59 ± 0.0002 mg GAE/g) than *E. intestinalis* extracts (41 ± 0.0002 mg GAE/g). In antimicrobial activity test, ethanol extracts *of H. musciformis* and *E. intestinalis* were found 10 mm of inhibition diameter against all of the bacterial strains. Besides, methanol extracts of *E. intestinalis* were more susceptible to *Staphylococcus aureus and Pseudomonas* which was close to the inhibition diameter (>15 mm) of the mainstream antibiotic; Gentamicin. Moreover, *Klebsiella sp*. was found more susceptible to ethanol and methanol extracts o*f Hypnea musciformis* as it showed inhibition zone greater than 15 mm. Both Seaweed extracts possessed higher amount of phytochemicals and showed promising antimicrobial activities when compared with the standards.

## 1. Introduction

Millions of people are being afflicted by different infectious diseases induced by pathogenic bacteria. Moreover, the phenomena of high mortality rate and the emergence of new multi-drug resistance, bacterial strains has become one of the threatening health problems worldwide [[1,2]. Different drug molecules are being applied to combat against those microorganisms. Antibiotics, metal ions, and various quaternary ammonium compounds are being used worldwide though these antimicrobial compounds are also being claimed to be associated with antibiotic resistance, complex chemical synthesis, environmental pollution, and high cost [3,4]. However, natural antibacterial agents have been recently identified to overcome these disadvantages [5]. Marine organisms are one of the richest sources of bioactive compounds and chemical diversity [6,7]. Different marine compounds, including seaweeds have historical contribution against the predators defense [8,9], and could be a promising antibacterial agents. [10,11].

Marine algae are one of the largest producers of biomass in the marine environmental [12]. Seaweeds are plant like ocean organisms that are botanically classified as microphysics marine algae. Edible seaweeds are often called “sea vegetables” seaweeds come in an amazing variety of beautiful shapes, colors and sizes and found in all of the world’s oceans. They are source materials for structurally unique natural products with pharmacological and biological activities [13]. Among the marine organisms, the macroalgae (seaweeds) occupy an important place as a source of biomedical compounds [14]. They are the most interesting algae group because of their broad spectrum of biological activities such as antimicrobial [15], antiviral [16], anti-allergic [17], anticoagulant [18], anticancer [19], antifouling [20] and antioxidant activities [21]. Importantly, seaweeds represent a potential source of antimicrobial substances due to their diversity of secondary metabolites with antiviral, antibacterial and antifungal activities [22]. Structurally diversed secondary metabolites of Seaweeds offer defense against herbivores, fouling organisms and pathogens and also play role in reproduction, protection from UV radiation and as allelopathic agents [23]. The bactericidal agents found in algae include aminoacids, terpenoids, phlorotannins, acrylic acid, phenolic compounds, steroids, halogenated ketones and alkanes, cyclic polysulphides and fatty acids [24]. Some of these metabolites extracted from seaweed such as iodine, carotene, glycerol, alginates, and carrageenans have been used in pharmaceutical industries [25,26].

The Bay of Bengal is the northeastern part of the Indian Ocean, bounded on the west and northwest by India on the north by Bangladesh. In Bangladesh, potential seaweeds are being reported in terms of food staffs and pharmaceutical agents from the south-eastern part of the mainland and offshore island. [27,28]. *Caulerparacemosa, Enteromorphasp, Gelidiellatenuissima, Gelidiumpusillum, Halymeniadiscoidea, Hypneapannosa, Hydroclathrusclathratus, Sargassumsp* are commercially important seaweeds in Bangladesh [29]. Moreover, settling a longstanding India-Bangladesh maritime boundary dispute, a Hague-based international court has recently awarded Bangladesh 19,467 square kilometers out of 25,602 sq km disputed area in the Bay of Bengal. Bangladesh is rich with around 133 species of seaweeds and eight of them are commercially important. Seaweed has great value in providing low-cost, wholesome nutrition and therapeutic protection. Still now, there are scanty information regarding the pharmaceutical potentiality of these species. That is why the objectives of the present study were to evaluate antimicrobial activity and phytochemical pattern of two widely available seaweed species (*Hypnea musciformis, Enteromorpha intestinalis*) of the Bay of Bengal sea coast.

## 2. Materials and Methods

### 2.1 Seaweed material and description of study area

*Two seaweeds species; H. musciformis* and *E. intestinalis* were used in this study. They were freshly collected from North Nuniar Chor, Cox’s bazar (21°35′0″N 92°01′0″E). (**Fig. 1**). Collected samples were washed in running water for 10 minutes and then transported to the laboratory and shade dried at 35±3 °C for 36 h. The shade dried seaweeds were powdered using electronic blender and used for further experiments.

**Fig. 1.**
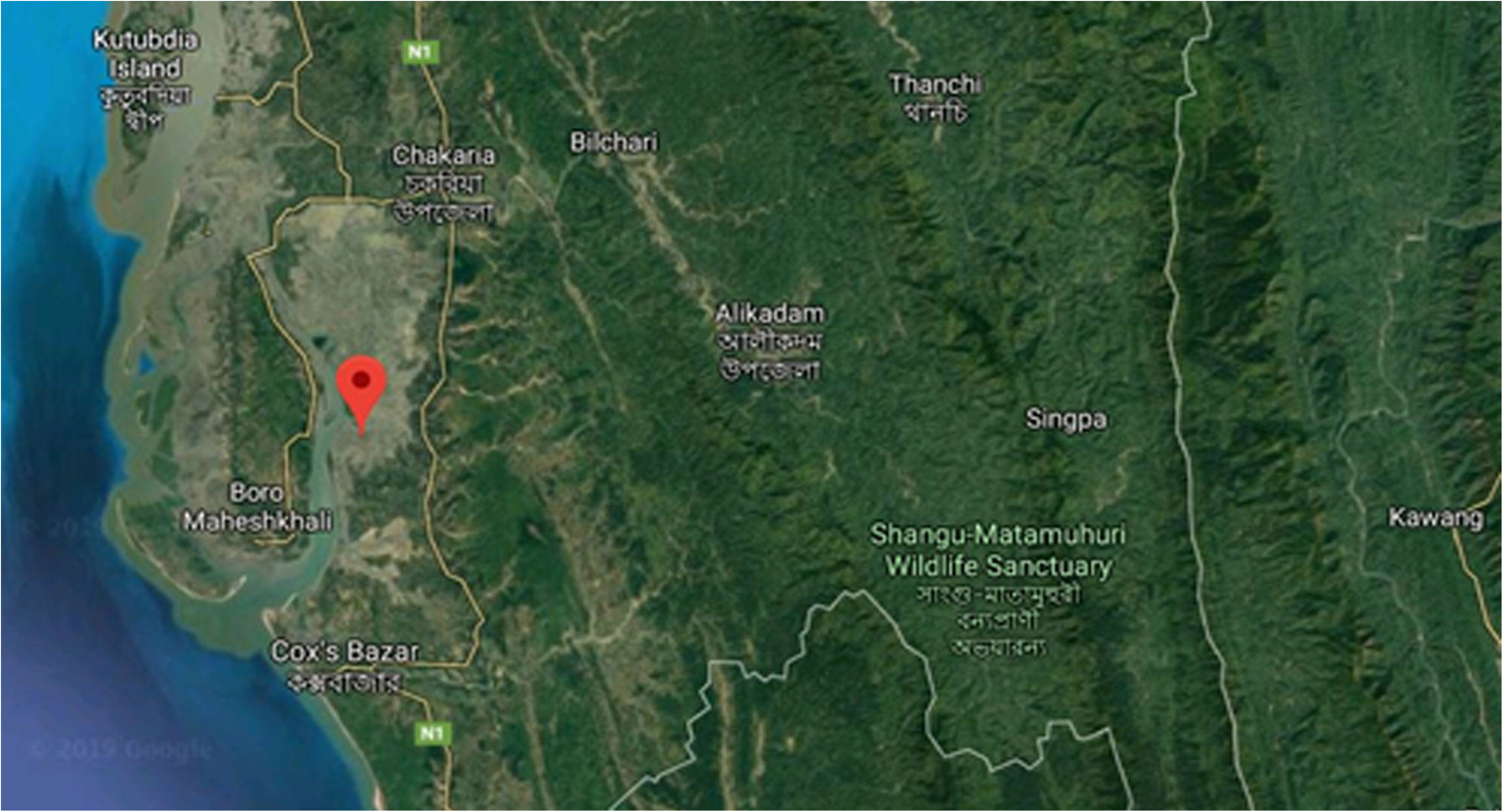
Sample collection site (21°35′0″N 92°01′0″E)

### 2.2 Preparation of seaweed extracts and fractions

The extracts were prepared by two different methods (water bath and soaking method). Each of 20 grams of powdered seaweed samples were soaked in a conical flask containing 160 mL of the distilled water, ethanol, methanol and acetone solvents. In soaking method, the samples were gently mixed by shaking and left for 72 hours at room temperature. In water bath method, the samples were mixed gently using the same solvents and left at 65°C for 4 hours. The liquid phase was then filtered with whatman no. 1 filter paper and allowed to be concentrated at reduced pressure to give specific solvent extracts. The crude extracts were used to test qualitative and quantitative analysis for secondary metabolites and antimicrobial activities against different bacterial strains.

### 2.3 Preliminary qualitative phytochemical analysis

Preliminary qualitative phytochemical analysis was carried out to identify the secondary metabolites present in the alcoholic extracts of *H. musciformis* and *E. intestinalis*.

#### 2.3.1 Flavonoids Test

For screening of flavonoids, sodium hydroxide test and shinoda test were conducted. After the addition of 2ml 10% aqueous sodium hydroxide solution, yellow color precipitation indicated the presence of flavonoid. Yellow color turned into colorless dilluted hydrochloric acid was added. In shinoda test, 2 drops of conc. HCl followed by a few fragments of magnesium ribbon were added. Formation of pink, reddish or brown color specifies the presence of flavonoids [30].

#### 2.3.2 Alkaloids Test

For this test, few drops of Wagner’s reagents or Dragendorff’s reagent in crude extracts were added. Appearance of a reddish-brown or orange red precipitation considered as the positive test for alkaloids [30,31,32,33].

#### 2.3.3 Test for Tannins

Addition of 2-3 drops of 5% ferric chloride to the crude extracts turned into brownish green or a blue-black color results the presence of tannin [32].

#### 2.3.4 Test for Phenols

A fresh mixture was produced from equal amount of 1% ferric chloride solution and 1% potassium ferrocyanide, 3 drops of the mixture added to the extract and filtered this solution, presence of phenol obtained from the formation of a bluish-green color [33].

#### 2.3.5 Test for Saponins

In a test tube, 2.5 ml of extract was added to 10 ml of sterile distilled water. The test tube was sealed with cap and shaken vigorously for about 30 seconds. It was then allowed to stand for 30 minutes. Formation of honeycomb froth indicated the presence of saponins [32].

### 2.4 Quantitative analysis of phytochemical constituency

Quantitative analysis for phytochemicals were performed for flavonoids, tannin, and phenolics

#### 2.4.1 Determination of Total Flavonoids Content

Total flavonoids content was determined by following the method by Wang [34]. About 0.2 mL of 10% AlCl_3_, 0.2mL of 1M potassium acetate and 5.6 mL of distilled water followed by 1 mL of methanolic extract at a concentration of 1 mg/mL was added in a test tube to estimate flavonoid content. Blank was prepared without methanol. A series of standard ((20, 40, 60,80, 100 μg/mL) was prepared using Quercetin followed by 0.2 mL of AlCl_3_, .2mL of 1M potassium acetate, 5.6 mL of distilled water. The mixtures were kept at room temperature for 30 minutes. Absorbance was taken at 415 nm using UV-vis spectrophotometer. The result was expressed in mg QE/g of the dried plant extractives.

#### 2.4.2 Determination of Total Tannins content

The tannins content was estimated by Folin-Ciocaltue’s method for both seaweed species as described by Kavita and Indira, 2016. [35] About 0.1 mL of the sample extract was added to a test tube containing 7.5 mL of distilled water and 0.5 mL of Folin-Ciocaltue phenol reagent. 1 mL of 35% Na_2_CO_3_ and dilute to 10 mL with distilled water. The mixtures were shaken well and kept at room temperature for 30 minutes. A set of reference standard solutions of tannic acid (20, 40, 60, 80, 100 μg/mL were prepared. Absorbance for test and standard solutions were measured against the blank at 700 nm with an UV/ Visible spectrophotometer. The estimation of the tannin content was carried out in triplicate. The tannin content was expressed in terms of mg of Tannic acid equivalent per g of dried sample.

#### 2.4.3 Determination of Total Phenolics content

The total phenolic content of dry extract was performed with Folin-Ciacaltue assay with slight described by Singleton et al., (1999) with slight modification [35,36]. About 1 mL of methanolic extract sample (1mg/mL) was mixed with 5 mL of 10% FolinCiaocalteu Reagent and 5 mL of 7.5% Na_2_CO_3_. Blank was concomitantly prepared, containing 1 ml methanol, 5 ml of 10% FolinCiocalteu’s reagent and 5 ml of 7.5% of NaHCO_3_. The mixtures were incubated for 20 minutes at 25°C followed by measuring absorbance taken at 760 nm against blank. Gallic acid was used as standard and reactions were performed as triplicates and mean value of absorbance was obtained. Calibration line was constructed using gallic acid as standard. The total content of phenol in extract was expressed in terms of Gallic acid equivalent (mg of GAE/g of extract).

### 2.5 Antibacterial activity of seaweeds extracts

#### 2.5.1 Microorganisms and media

Five bacteria species obtained from the Laboratory of Microbiology, Department of Genetic Engineering and Biotechnology, Shahjalal University of Science and Technology and used as the antimicrobial test strains: There were four gram negative (*Escherichia coli, Klebsiella sp., Pseudomonas sp., Salmonella sp*.) and one gram positive (*Staphylococcus aureus*) bacteria. The bacterial strains were maintained on the nutrient agar medium [36].

#### 2.5.2 Agar disk-diffusion assay

The screening of antimicrobial activity of the seaweed extracts was carried out with agar disk-diffusion method using Muller Hinton Agar (MHA) medium [31,37]. Bacterial culture (50 μL) was taken from the nutrient broth culture using 100 μL micropipette and poured into the sterile plate containing Muller-Hinton agar medium. Sterile cotton was used for streaking the dried surface of plates. Under aseptic condition, prepared discs (5 mm round filter paper soaked with test solution at a concentration of 1mg/mL) were air dried, placed into center of an agar plate by using a sterile forceps and pressed down. Then discs were employed to be incubated at 37°C within 15 minutes. After 24 & 48 hours of incubation, each plate was examined. There were uniformly circular zone of inhibition on the surface. The diameter of the complete zone of inhibition was measured. All tests were performed in triplicate manner.

## 3. Results

### 3.1 Preliminary qualitative phytochemical analysis

The present study exposed that ethanolic extracts of *H. musciformis* and *E. intestinalis* contained different plant secondary metabolites *viz* alkaloids, flavonoids, tannins, phenols, and saponins. Preliminary phytochemicals study revealed that *E. intestinalis* extracts contained flavonoids and saponins in higher extent on the other hand alkaloids and tannins were present in *Hypnea musciformis* in greater content. Other secondary metabolites were moderately present in two seaweed extracts (**Table 1**).

**Table 1.** Phytochemical Screening of *Hypnea musciformis* and *Enteromorpha intestinalis*

| *Name of Test* | *Method of Test* | *Seaweed* |  |
| --- | --- | --- | --- |
|  |  | *Hypnea musciformis* | *Enteromorpha intestinalis* |
| Flavonoids | Sodium Hydroxide test | ++ | ++ |
|  | Shinodas test | ++ | +++ |
| Alkaloids | Wagner's test | +++ | + |
|  | Dragendorff's test | ++ | + |
| Tannins | Potassium dichromate test | +++ | ++ |
|  | Ferric chloride test | ++ | ++ |
| Phenols | - | ++ | ++ |
| Saponins | Froth test | ++ | +++ |

[+++ : Higher presence, ++ : Moderate presence, ++ : Low presence]

### 3.2 Quantitative determination of the phytochemical constituency

**Table 2** showed the qualitative determination of the tannin, flavonoid and phenolic content. Folin-Ciocaltue’s method was used to determine the total tannin content in different seaweed extracts. The tannin content was expressed in terms of mg of Tannic acid equivalent per g of dried sample. Ethanolic extracts of *H. musciformis and E. intestinalis* estimated 51 ± 0.0002 and 43 ± 0.0002 mg/g respectively. However, total flavonoid contents of the extracts were expressed in terms of quercetin equivalent (QE). *H. musciformis* and *E. intestinalis* estimated 67 ± 0.0002 and 57 ± 0.0002 mg/g respectively. Total phenolic contents of the extracts were determined in terms of gallic acid equivalent (GAE). *H. musciformis* exhibited higher amount of phenolics (59 ± 0.0002) present in the crude extracts rather than *E. intestinalis* extracts (41 ± 0.0002).

**Table 2.**
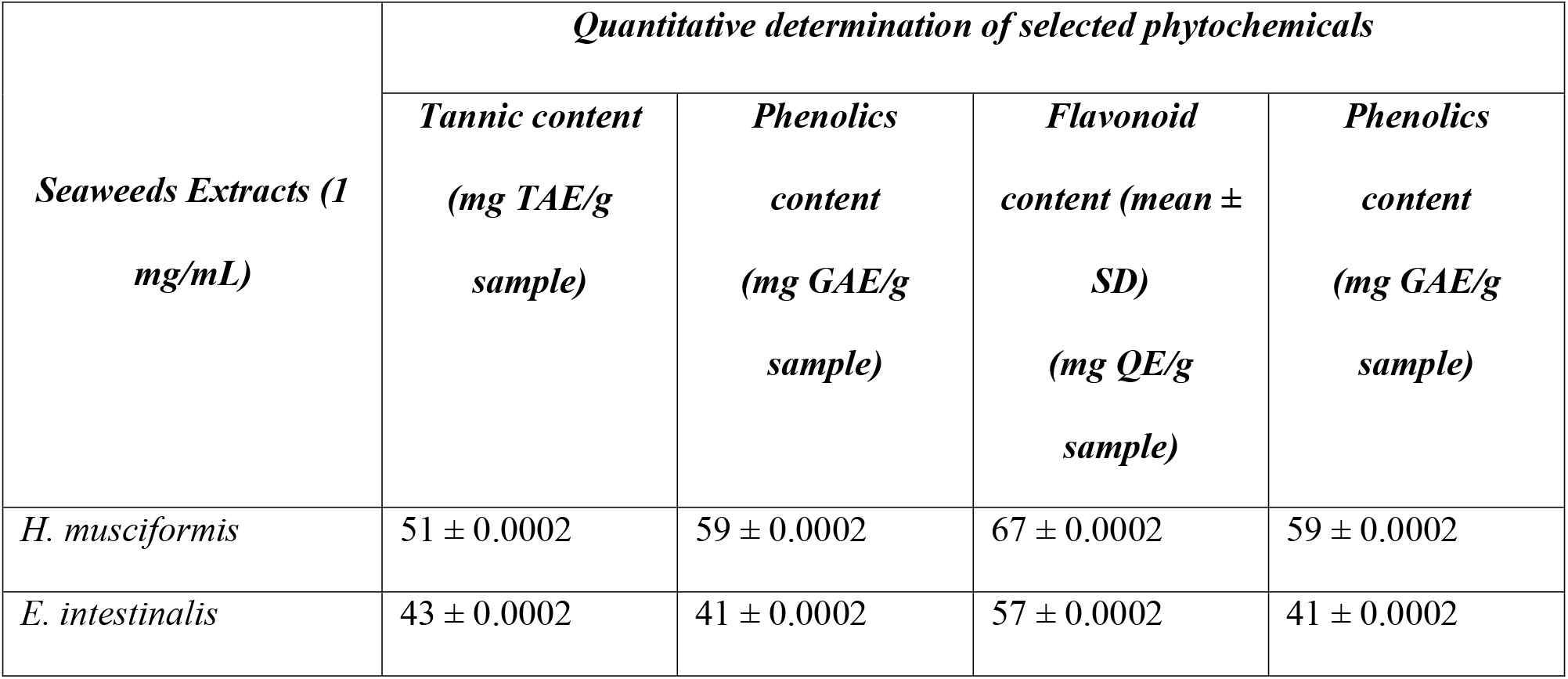
Quantitative determination of phytochemical constituency

### 3.3 Antibacterial activity of seaweeds extracts

Two extraction methods, for instances, soaking and water bath method were applied for generating seaweeds extracts by using ethanol, methanol, acetone and distilled water. The results of antibacterial activities of the seaweeds extracts against selected bacterial strains are summarized in **Table 3** in terms of socking method and **Table 4** for water batch method.

**Table 3.** Screening of antibacterial activity of seaweed extracts by soaking method

| *Seaweed* | *Solvent* | *Gram positive*<br>*bacteria* | *Gram negative bacteria* |  |  |  |
| --- | --- | --- | --- | --- | --- | --- |
|  |  | *S. aureus* | *E.coli* | *Klebsiella*<br>*sp.* | *Pseudomonas*<br>*sp.* | *Salmonella*<br>*sp.* |
| *Hypnea*<br>*musciformis* | Ethanol | ++ | ++ | ++ | ++ | ++ |
|  | Methanol | + | ++ | + | ++ | ++ |
|  | Acetone | - | + | + | + | + |
|  | Distilled<br>water | ++ | ++ | ++ | ++ | ++ |
| *Enteromorpha*<br>*intestinalis* | Ethanol | ++ | ++ | ++ | ++ | ++ |
|  | Methanol | +++ | ++ | ++ | +++ | ++ |
|  | Acetone | +++ | + | ++ | ++ | ++ |
|  | Distilled<br>water | ++ | + | ++ | ++ | ++ |
| Gentamicin<br>(Control) |  | +++ | +++ | +++ | +++ | +++ |

—: no activity; +: inhibition diameter < 10 mm; ++: inhibition diameter < 15 mm; +++: inhibition diameter < 20 mm;

**Table 4.** Screening of antibacterial activity of seaweed extracts by water bath method

| *Seaweed* | *Solvent* | *Gram positive*<br>*bacteria* | *Gram negative bacteria* |  |  |  |
| --- | --- | --- | --- | --- | --- | --- |
|  |  | *Staphylococcus*<br>*aureus* | *E. coli* | *Klebsiella*<br>*sp.* | *Pseudomonas*<br>*sp.* | *Salmonella*<br>*sp.* |
| *H. musciformis* | Ethanol | ++ | ++ | +++ | ++ | ++ |
|  | Methanol | ++ | ++ | +++ | ++ | ++ |
|  | Acetone | ++ | ++ | ++ | ++ | ++ |
|  | Distilled<br>water | ++ | ++ | ++ | ++ | ++ |
| *E. intestinalis* | Ethanol | ++ | ++ | ++ | ++ | ++ |
|  | Methanol | ++ | ++ | ++ | ++ | + |
|  | Acetone | ++ | ++ | ++ | ++ | ++ |
|  | Distilled<br>water | + | ++ | ++ | ++ | ++ |
| Gentamicin<br>(Control) |  | +++ | +++ | +++ | +++ | +++ |

—: no activity; +: inhibition diameter < 10 mm; ++: inhibition diameter < 15 mm; +++: inhibition diameter < 20 mm;

In case of soaking method, ethanol extracts of *H. musciformis* and *E. intestinalis* were found more than 10nm inhibition zone against all of the bacterial strains. But, methanol extracts of *E. intestinalis* were found more active against *Staphylococcus aureus Pseudomonas* as these showed inhibition diameter greater than 15 mm, which was similar to our studied control; Gentamicin (**Fig. 2**). In soaking method ethanol extract of *Hypnea musciformis* showed highest zone of inhibition (14 ± 0.76 mm) against *E. coli* and methanol extract of *E. intestinalis* showed highest zone of inhibition against *S. aureus* (**Fig. 2**). Acetone extract of *H. musciformis* show highest zone of inhibition against *S. aureus* but which is less than *E.intestinalis*.

**Fig. 2.**
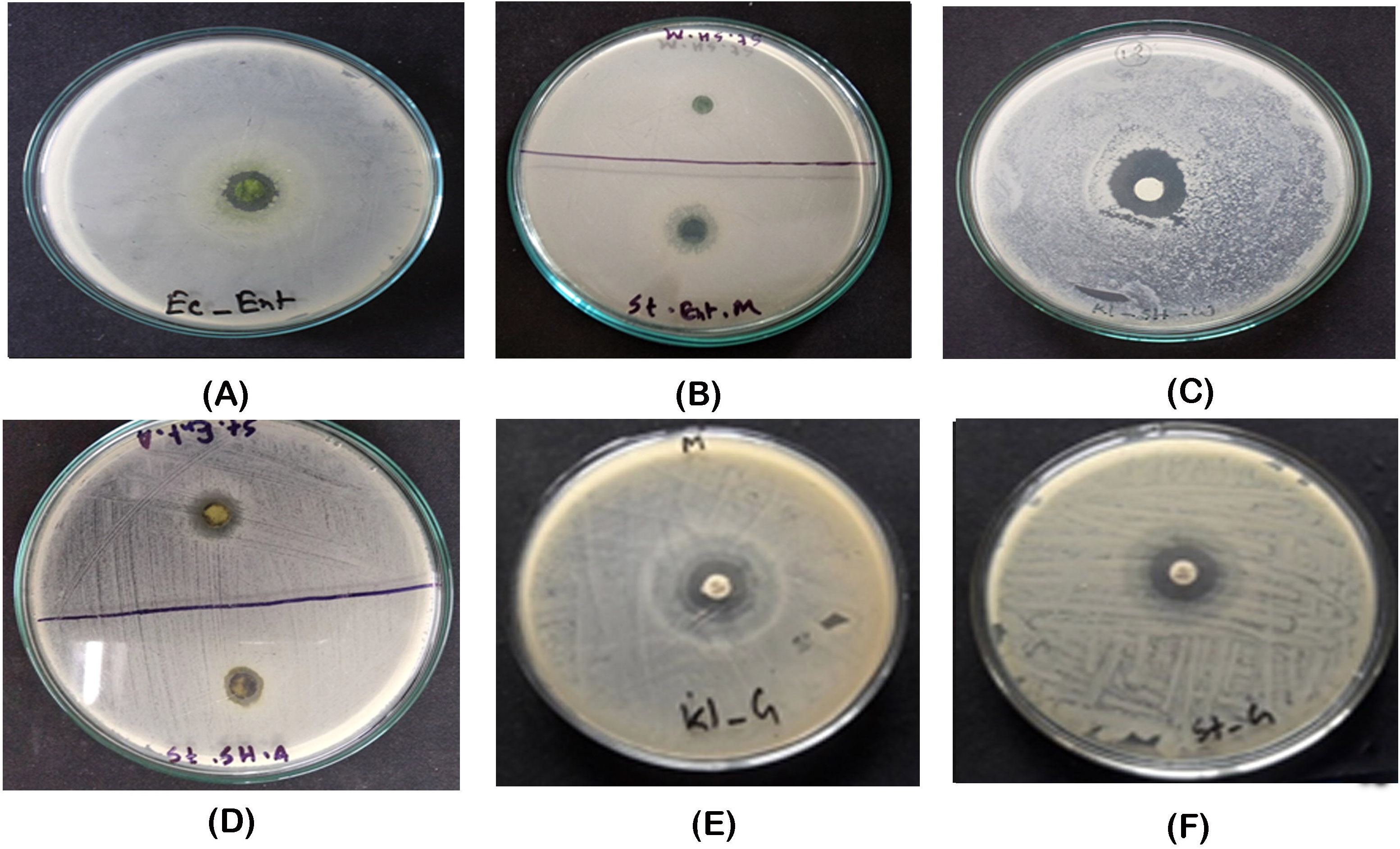
Growth inhibition zone of of different extracts of *Hypnea musciformis* and *Enteromorpha intestinalis*. (A) *E. intestinalis* ethanol extract against *E. coli* (soaking method). (B) *E. intestinalis* and *H. musciformis* methanol extract (soaking method) against *S. aureus*. (C) *H. musciformis* ethanol extract against *Klebsiella sp*. (Water bath method). (D) *E. intestinalis* and *H. musciformis* acetone extracts against *S. aureus*(soaking method). (E) Gentamicin against *Klebsiella sp*. (F) Gentamicin against *Staphylococcus aureus*.

In case of water bath method, *Klebsiella sp*. was found more susceptible to ethanol and methanol extracts of *Hypnea musciformis* as it showed inhibition zone greater than 15 mm (**Fig. 2**). However, in terms of *Enteromorpha intestinalis*, inhibition zone was found more than 10nm in maximum antimicrobial tests against all bacterial strains. Again, *H. musciformis* exhibited more active zone of inhibition against different microbes rather than *E. intestinalis*.

## 4. Discussion

Phytochemical profiling of the seaweed samples revealed the presence of different phytochemicals in *Hypnea musciformis* and *Enteromorpha intestinalis*. Phenol, tannin and flavonoids were found in greater amount in both extracts [38,39]. These phytochemicals could exhibit antimicrobial activity for presence of the phytochemical constituency. Rojas *et al*. indicated that the antibacterial activity is due to different chemical agents present in the extract, including flavonoids and triterpenoids and other compounds of phenolic nature or free hydroxyl group, classified as active antimicrobial compounds [40].

Variation in antibacterial activity may be due to the method of extraction, solvent used in extraction and season at which samples were collected [37]. Several different organic solvents have been used to screen algae for antibacterial activity [41]. In current study we used alcoholic and aqueous solvents to generate seaweeds extracts. Ethanol extract of *E. intestinalis* showed highest zone of inhibition against *E. coli*. and methanol extract of *E. intestinalis* showed highest zone of inhibition against *S. aureus*. Ethanol extract of *Enteromorpha intestinalis* and methanol extracts of *Hypnea musciformis* were more effective against selected bacterial strains.

The both seaweed extracts (*E. intestinalis, H. musciformis*) are effective against gram positive and gram negative bacteria. Kolanjinathan K, Stella D showed that *Ulva lactua, Halimedagracilis, Gracilaria edulis, Hypnea musciformis, Turbinariaconoides, Sargassum myricystum* effective against *E. coli, P. aeruginosa, S. aureus, K. pneumoniae, E.faecalis* [26,42] which supports the study results. There are some other studies which could strengthen our findings, for example, Tuney I *et al*. illustrated that *Enteromorpha sp*. is *effective* against *Candida* sp., *E. faecalis, S. aureus, S. epidermidis, P. aeruginosa, E. coli* [43]. Sukatar A *et al*. reported that *Enteromorpha linza* is highly suspectible to *S. aureus, S. epidermidis, S. fecalis, B. subtilis, S. typhimurium, P. aeruginosa, E. cloacae, E. coli, C. albicans* [44]. Two different extraction methods were used. Among them *Hypnea musciformis* showed better results in water bath extraction method and *Enteromorpha intestinalis was better* in soaking method. So it could be assumed that *Hypnea musciformis* release more bioactive compound in water bath method and *Enteromorpha intestinalis* in soaking method.

## 5. Conclusion

It could be concluded that the both studied seaweeds, *Hypnea musciformis* and *Enteromorpha intestinalis* possessed potential antimicrobial activities. It may be due the availability of secondary metabolites which could induce antimicrobial approaches. The strong correlation between the contents of different secondary metabolites, such as phenols, tannins and flavonoids employs that these phytochemicals and prime contributors to the antimicrobial potentiality of these seaweeds species.

## Acknowledgment

We are highly thankful to Professor Dr Abul Kalam Azad as he provided us the pathogenic bacterial strains for the study.

